# Stochasticity contributes to explaining minority and majority MOMP during apoptosis

**DOI:** 10.1101/2025.05.02.651813

**Authors:** Jenny Geiger, Fabian Klötzer, Nadine Pollak, Gavin Fullstone, Markus Rehm

## Abstract

Apoptosis dysfunction is linked to diseases like cancer and neurodegenerative disorders. A key event during apoptosis is mitochondrial outer membrane permeabilization (MOMP), which typically proceeds in a rapid all-or-none fashion. If MOMP occurs only in a subset of mitochondria (minority MOMP), it can be sublethal and contribute to tumorigenesis and cancer progression. Similarly, individual mitochondria escaping widespread MOMP (majority MOMP) can allow cancer cells to recover if apoptosis execution fails. How such heterogeneities in mitochondrial MOMP responsiveness arise within cells is incompletely understood. In particular, whether stochasticity in subcellular protein distributions and interactions across cytosol and mitochondria can realistically contribute to mitochondrial MOMP heterogeneity has not yet been studied. To assess this, we sequentially built and experimentally parameterized a particle-based, cell-sized model including cytosolic and mitochondrial compartments, and that featured a reduced interactome of MCL-1, BAK and tBID. High-performance computing enabled cell-scale simulations of protein distributions and interactions to understand how and under which conditions stochasticity could contribute to heterogeneity in MOMP susceptibility. Our results show that stochastic effects strongly predispose sub-pools of fragmented mitochondria to MOMP under low apoptotic stress. At higher apoptotic stress, fractions of small mitochondria were more likely to escape MOMP than large mitochondria. Retrospective quantification of mitochondrial sizes in experimental scenarios of minority and majority MOMP confirmed these findings. We therefore conclude that stochasticity substantially contributes to enabling small or fragmented mitochondria to undergo MOMP in minority MOMP scenarios and to escape MOMP in majority MOMP scenarios.

## Introduction

Apoptotic cell death plays an essential role in development, tissue homeostasis, and immune regulation by ensuring that damaged or unnecessary cells are eliminated. When dysregulated, it can contribute to various diseases, including cancer, autoimmune disorders, and neurodegeneration (1).

Mitochondria play a central role in the process of apoptotic decision making. During apoptosis, mitochondria release pro-apoptotic factors such as cytochrome-c and Smac into the cytosol, initiating the caspase-dependent execution phase of cell death (2–4). The release of these factors is controlled by the formation of pores in the mitochondrial outer membrane, a process relying on pro-apoptotic multi-domain members of the BCL-2 protein family, such as BAK and BAX (5). BAK resides predominantly in the mitochondrial outer membrane, whereas BAX typically is predominantly found in the cytosol and translocates to the mitochondria in response to apoptotic stress. The activation of BAK and BAX is triggered by BH3-only proteins, a subclass of the BCL-2 protein family that includes proteins such as BID/tBID and PUMA (1,6,7). Anti-apoptotic BCL-2 family proteins, such as MCL-1, BCL-XL, and BCL-2 itself (8–11), instead act as safeguards against mitochondrial outer membrane permeabilization (MOMP). These proteins prevent pore formation by binding and neutralizing BH3-only proteins as well as activated BAK and BAX (1,12).

In most scenarios, all mitochondria undergo MOMP, leading to a rapid and kinetically largely invariant release of cytochrome-c and Smac, and, within minutes, a swift activation of effector caspases (13–15). However, scenarios exist in which MOMP does not occur within all mitochondria. For example, treatments with BH3-mimetics, if not lethal, can trigger MOMP in only a fraction of the overall pool of mitochondria (minority MOMP) (16). Similarly, if cells incapable of activating effector caspases undergo widespread MOMP, a few mitochondria might escape MOMP and can contribute to cell recovery (17).

The reasons for heterogeneities in MOMP susceptibility between mitochondria within the same cell remain incompletely understood. However, various studies have proposed or demonstrated that features which in principle can differ between mitochondria, and which can be measured experimentally, could contribute to inter-mitochondrial differences in MOMP susceptibility. Among these are, for example, mitochondrial bioenergetics and metabolic fitness, membrane curvature, membrane lipid composition, but also mitochondrial proteins that could support pore formation by BAK and BAX (18–24). What remains inaccessible by experimental observation of cells and their mitochondrial pools is if stochasticity in the combined movement and interaction of multiple MOMP-regulatory proteins could in principle and realistically also contribute to inter-mitochondrial heterogeneities in MOMP susceptibility. However, simulation-based approaches can be suitable to provide such estimates and potentially link these to observed phenotypes. The development of a theoretical framework implying that the stochastic movement (“random walk”) of molecules in solution underlies the macroscopic observation of diffusion and Brownian motion dates back more than a century (25–27). In prior work, we could describe diffusion-related phenotypes during apoptosis signal transduction by deterministic spatiotemporal modelling at cell scale (14,28). However, stochastic modelling of MOMP regulation had to be limited to simulations of small patches of mitochondrial membrane areas (29), owing to the high computational demand of such simulations. With access to modern high-performance computing, we now were able to develop an experimentally parameterized cell-scale and particle-resolution model of a reduced MCL-1/BAK/tBID interactome. We used this model to estimate if and under which conditions stochastic effects could play a role in giving rise to heterogeneities in MOMP susceptibility, and to study if modelling results align with experimental observations by retrospective data analysis.

## Results

To assess the influence of stochasticity on MOMP regulation, the most powerful suitable simulation approaches are particle-based models that capture the heterogeneity and stochastic features of protein behaviours in solution and within membranes. To simulate the spatiotemporal distribution of BCL-2 family proteins in such a particle-based model, we first aimed to establish a simplified *in silica* 3D cell model comprising of mitochondria and the cytosol. This model was then used to study if a representative BCL-2 family member, in this case MCL-1, behaves as expected, and which influence mitochondrial arrangements and sizes have on subcellular protein distributions. This model was then expanded with BAK as a representative pore forming protein, and tBID as an activator of BAK and inhibitor of MCL-1, to allow for the simulation of BAK oligomerization at single mitochondrion resolution.

### A fundamental particle-based model shows that MCL-1 collisions with mitochondria correlate with mitochondrial surface areas

In the initial particle-based model, we implemented MCL-1 as dynamically (retro)translocating between the cytosol and mitochondria (30) (**Fig. 1A**). For the parameterization of cell volume and MCL-1 amounts, in-house experimental data from NCI-H460 cells, a widely used lung cancer cell line (31), were generated (**Supplemental Fig. 1A**, **B**). (Retro)translocation dynamics of MCL-1 were implemented based on imaging data-derived rates from these cells (see methods). The resulting model accurately reflected the mean steady-state ratio between cytosolic and mitochondrial MCL-1 (visualization in **Fig. 1B**; **Fig. 1C**). On purpose, translocation/retrotranslocation rates were not assumed to vary in the model, so that heterogeneities arising in subsequent simulation-based experiments could clearly and causally be linked to respective parameter changes. Cytosolic MCL-1 particles were implemented to diffuse in 3D in the cytosol, whereas the diffusion of mitochondrial MCL-1 was assumed as lateral random walk within the outer mitochondrial membrane. Mitochondrial networks are dynamically remodelled by fission and fusion events, e.g. to ensure a sufficient number of mitochondria in daughter cells after mitosis and to allow mitochondria to be distributed and transported throughout large, outstretched cells, such as neurons (32–34). As a consequence of such dynamics, mitochondrial surface areas keep changing and can affect interactions with mobile proteins. To test if changes in the mitochondrial surface area influence the subcellular distribution of MCL-1 as expected, we changed the mitochondrial surface area in the model cell while trying to keep the total mitochondrial volume constant (illustrated in **Fig. 1D**). Indeed, the interactions (collision rates) of MCL-1 particles with the mitochondrial membrane correlated proportionally with the mitochondrial surface area (**Fig. 1E**). As a consequence of this, the ratio of cytoplasmic MCL-1 (MCL-1_Cyto_) to mitochondrial MCL-1 (MCL-1_Mito_) amounts correlated inversely exponentially to the mitochondrial surface area (**Fig. 1F**). Our particle-based model of a simplified three-dimensional cell therefore sufficed to replicate subcellular MCL-1 distributions and satisfied expected protein behaviours upon perturbing mitochondrial surface areas. Based on this implementation, we next studied the influence of sub-cellular mitochondrial arrangements on the heterogeneity of MCL-1 distributions.

**Figure 1:**
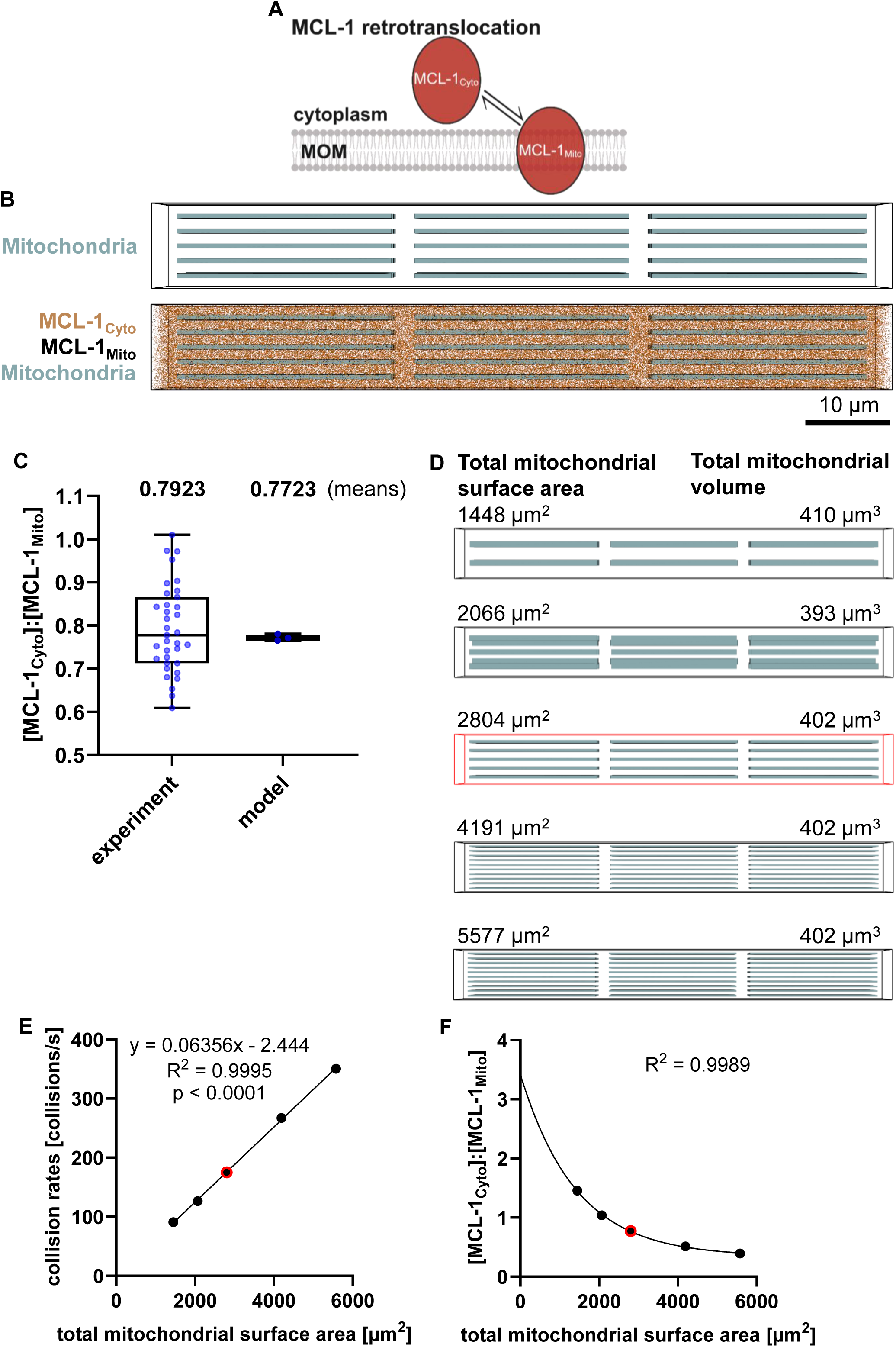
Particle-based modelling of MCL-1 localization reveals a correlation between mitochondrial surface area and mitochondrial MCL-1 accumulation. (A) Schematic overview of MCL-1 retro-translocation between the mitochondrial outer membrane (MOM) and the cytoplasm. (B) Visualization of the simplified model cell at steady state. Mitochondria are shown in blue, cytosolic MCL-1 in orange, and mitochondria-bound MCL-1 in black. (C) The model reproduces an experimentally observed steady-state MCL-1 distribution. The model results are based on three independent simulation runs with randomized initial particle/MCL-1_Cyto_ distributions. (D) Visualization of model cells with different mitochondrial surface areas, while keeping total mitochondrial volume relatively constant. The model cell used in B-C is highlighted in red. (E) MCL-1-mitochondria collision rates were quantified. Results represent the means of three simulation runs with varying initial particle distributions. Variances were too small to be displayed. (F) Subcellular MCL-1 distributions of simulations with (retro-)translocation were quantified A one-phase decay fit was applied. Results represent the means of three simulation runs with varying initial particle distributions. Variances were too small to be displayed.

### Asymmetric mitochondrial positioning gives rise to heterogeneities in the amounts of mitochondrial MCL-1

To eliminate mitochondrial positioning as a variable, we had constructed the initial model by regularly and symmetrically arranging mitochondria of the same size (**Fig. 1B**). However, mitochondria are not uniformly distributed across cells, but cells instead can present with areas enriched or deprived of mitochondria (35,36). Furthermore, active mitochondrial trafficking dynamically changes mitochondrial positions and densities throughout a cell’s lifetime (33). To assess the influence of mitochondrial positioning on cellular MCL-1 distributions, we randomized the positions of the mitochondria within the model cell (model visualization in **Fig. 2A**). The randomized positioning did not affect median particle densities of mitochondrial MCL-1 when related to mitochondrial volume or surface area (**Fig. 2B, C**). However, the random arrangement significantly increased the variances in particle densities (**Fig. 2B, C**). To ensure the robustness of this finding and to exclude potential skewing of the model performance due to the geometry of the model cell, we validated this observation further. Hereby, we set up a series of extreme cases of mitochondrial positioning within the model cell, where the mitochondria were randomly seeded exclusively at the top and bottom, front and back, or left and right sides of the cell (illustrated in **Fig. 2D**). The median of mitochondrial MCL-1 particles was comparable between all scenarios, and likewise the variances of MCL-1 densities remained comparable between these random positioning scenarios (**Fig. 2E, F**). As such, random mitochondrial arrangements, including dense and sparse regions, can contribute to inter-mitochondrial heterogeneity in MCL-1 concentrations.

**Figure 2:**
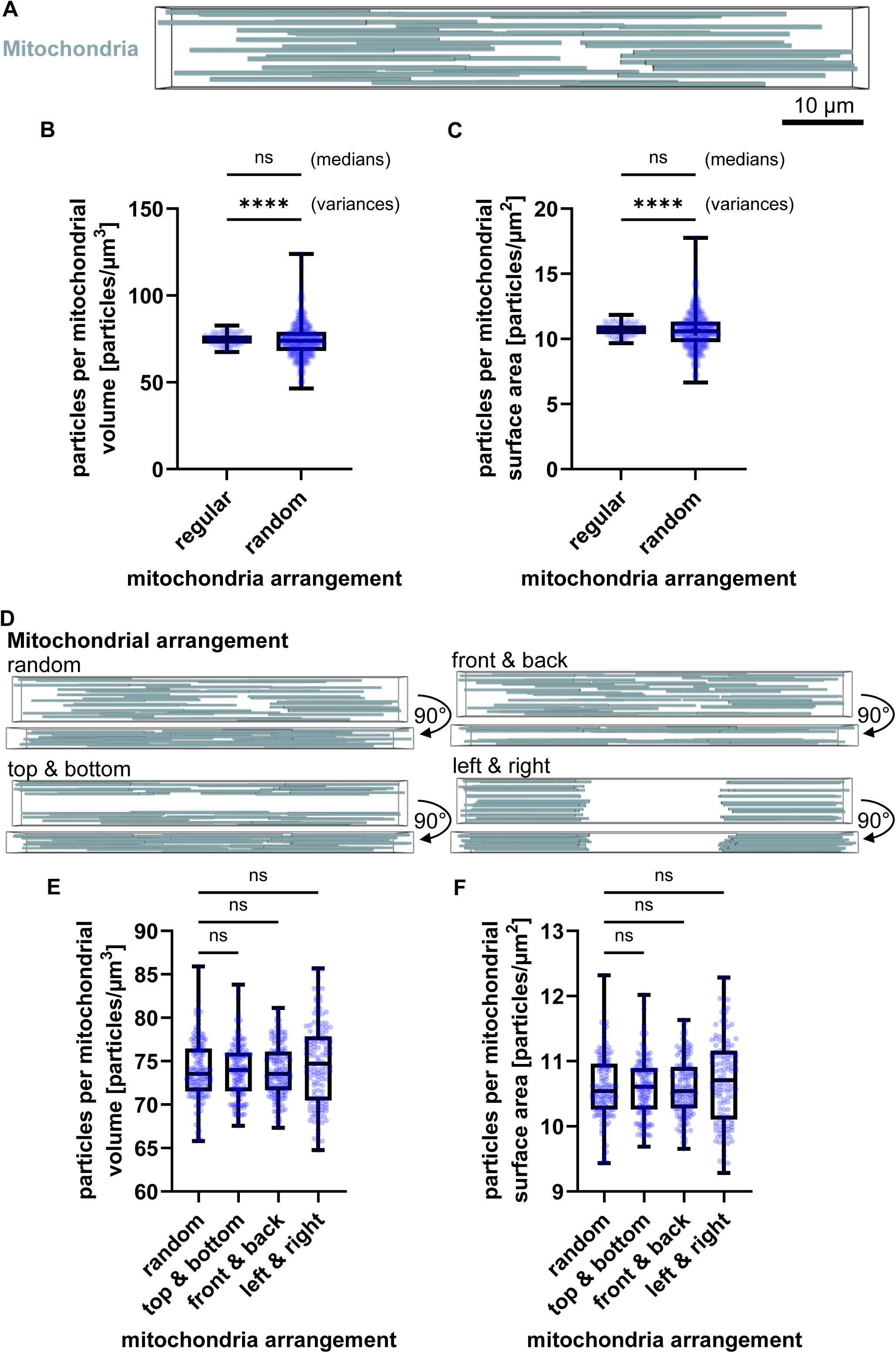
Asymmetric mitochondrial positioning gives rise to heterogeneities in the amounts of mitochondrial MCL-1. (A) Visualization of a model cell with mitochondria (blue) randomly distributed throughout the cytoplasm. (B-C) Quantification of mitochondrial MCL-1 density per mitochondrial volume (B) and per surface area (C), comparing regular and random mitochondrial arrangements. Medians comparison by Mann-Whitney U test: (E) p = 0.1347 and (F) p = 0.1333. Variances comparison by Levene’s test (p < 0.0001). (D) Model visualization for three extreme mitochondrial arrangements (top/bottom, front/back, left/right). Upper panels show frontal view; lower panels show top-down view. MCL-1 particles are hidden for clarity. (E-F) Quantification of mitochondrial MCL-1 particles per volume (E) and per surface area (F) across the extreme spatial arrangements shown in (D). Medians comparison by Kruskal-Wallis test followed by Dunn’s post-hoc test for multiple comparisons: (E) p = 0.2945 with p ≫ 0.05 for all multiple comparisons and (F) p = 0.293 with p ≫ 0.05 for all multiple comparisons.

### Mitochondrial fission and fragmentation increase the variance of mitochondrial-bound MCL-1 particles

The mitochondrial network is a highly dynamic system that undergoes continuous remodelling through fission and fusion events, which serve as quality control mechanisms to maintain cellular homeostasis (37,38). Cellular stress conditions can impair mitochondrial fitness and cause mitochondrial fragmentation, while increased cellular energy demands and cellular responses to counteract stress conditions can promote mitochondrial fusion and cell survival (37,38).

Therefore, we next investigated if and to what extent mitochondrial fission and fragmentation could be a cause for inter-mitochondrial heterogeneities in MCL-1 amounts to arise as a result of stochasticity in particle-scale diffusion. To this end, we replaced 10% of the mitochondria with shorter, small mitochondria (visualization in **Fig. 3A**), while neither the total mitochondrial volume nor the kinetic parameters were modified (**Supplemental Fig. 3A**). MCL-1 particle concentrations did not differ in non-fragmented mitochondria between the original model and the model variant including mitochondrial fragmentation (**Fig. 3B**). Since fragmented mitochondrial have a higher surface-to-volume ratio, we expected a higher number of mitochondrial-bound particles per mitochondrial volume specifically in fragmented mitochondria. Indeed, our simulations confirmed such an increase for fragmented mitochondria (**Fig. 3B**). Since protein interactions relevant for the regulation of MOMP occur at or within the outer mitochondrial membrane, it is more relevant to estimate the impact of fragmentation on MCL-1 density, i.e., MCL-1 particles per mitochondrial surface area. While the median number of mitochondrial-bound particles per mitochondrial surface area remained comparable between fragmented and non-fragmented mitochondria, fragmentation resulted in a significantly higher variance in mitochondrial-bound particle densities (**Fig. 3C**). This finding is particularly relevant because it suggests that mitochondrial fragmentation does not simply lead to a uniform increase or decrease in particle densities. Instead, at the scale of fragmented mitochondria, heterogeneities can emerge due to the stochastic random walk and (retro)translocation of diffusing MCL-1 particles at realistic intracellular concentrations. As a consequence, distinct subpopulations of mitochondria may form, with either high or low MCL-1 densities.

**Figure 3:**
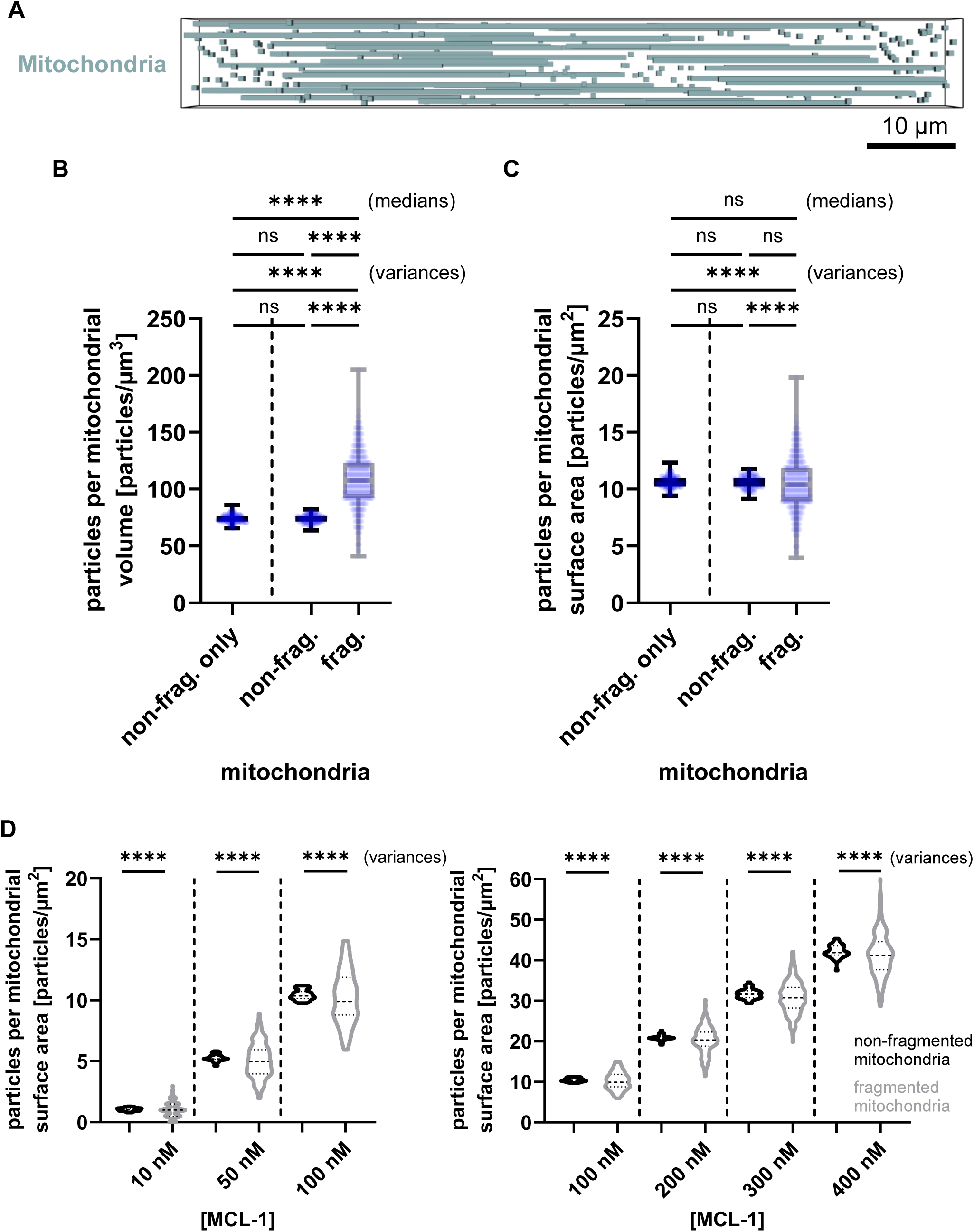
Mitochondrial fragmentation increases the variance of mitochondrial-bound particles. (A) Visualization of the particle-based model with implementation of mitochondrial fragmentation (10% of mitochondria). Total mitochondrial volume was kept constant. Mitochondria are shown in blue; particles are hidden for clarity. (B) Quantification of mitochondrial MCL-1 per mitochondrial volume in non-fragmented mitochondria only (from Fig. 2A), as well as in cells with non-fragmented and fragmented mitochondria (shown in A). Medians comparison by Kruskal-Wallis test followed by Dunn’s post-hoc test (p > 0.05 between non-fragmented mitochondria and p < 0.0001 between non-fragmented and fragmented mitochondria). Variances comparison by Brown-Forsythe test (p > 0.05 between non-fragmented mitochondria and p < 0.0001 between non-fragmented and fragmented mitochondria). (C) Quantification of mitochondrial MCL-1 per mitochondrial surface area. Medians comparison by Kruskal-Wallis followed by Dunn’s post-hoc test (p > 0.05). Variances comparison by Brown-Forsythe test (p > 0.05 between non-fragmented mitochondria and p < 0.0001 between non-fragmented and fragmented mitochondria). D’Agostino-Pearson tests confirmed normal distribution in all groups. (D) Analysis of mitochondrial MCL-1 per mitochondrial surface area across a range of initial MCL-1 concentrations (10–400 nM). Medians comparison by Kruskal-Wallis (p > 0.05). Variances comparison by Brown-Forsythe (p < 0.0001). Datapoints represent individual mitochondria pooled from 3 separate simulations of a single cell.

We next studied if this stochasticity-driven heterogeneity on fragmented mitochondria is maintained also at MCL-1 concentrations lower or higher than the reference total MCL-1 concentration of 101 nM. To test this, we varied MCL-1 concentrations from 10 nM to 400 nM. Comparative analysis revealed that fragmented mitochondria always exhibited significantly greater variance in mitochondrial MCL-1 particles per surface area than non-fragmented mitochondria (**Fig. 3D**), and this finding was robustly observed in independent repeat simulations (**Supplemental Fig. 3B, C**). It can be assumed that this finding extends to other members of the BCL-2 family, both pro- and anti-apoptotic. Consequently, such variabilities in distributions may contribute to differences in MOMP susceptibility between non-fragmented and fragmented mitochondria and also to differential inter-mitochondrial responses under stress conditions.

### Mitochondrial fragmentation increases variability in MOMP susceptibility due to stochasticity in protein distributions

Heterogeneity between fragmented mitochondria due to stochastic effects on protein distributions including and beyond MCL-1 might give rise to heterogeneous MOMP susceptibilities in these mitochondrial subpopulations. To investigate this, the particle-based model was expanded to include additional members of the BCL-2 protein family. Specifically, we developed a minimal apoptosis system, similar to previous deterministic modelling of MOMP susceptibilities (39), by including the pro-apoptotic pore-forming protein BAK and the BH3-only protein tBID.

BAK was implemented with an experimentally determined concentration (**Supplemental Fig. 1A**). tBID, which is produced in cells upon receiving extrinsic apoptosis signals, was initially implemented at a concentration of 10 nM, resembling a strong pro-apoptotic signal (40). When seeding simulated cells with these proteins, BAK and tBID particles per area strongly varied between fragmented mitochondria, similar to MCL-1 (**Fig. 4 A-C**). We implemented tBID as an inhibitor of MCL-1 and as an activator of BAK, and MCL-1 as an inhibitor of activated BAK (aBAK) and, accordingly, as a neutralizer of tBID. Activated BAK was assumed to form multiples of dimers (41), with tetra- and hexamers assumed to be sufficient for forming pores in the outer mitochondrial membrane (42). The parameterization was derived from a prior deterministic modelling approach (39) by converting forward and reverse interaction rates to parameters suitable for stochastic particle-based modelling (**Fig. 4D, Supplemental Fig. 1E**, **F and Methods**).

**Figure 4:**
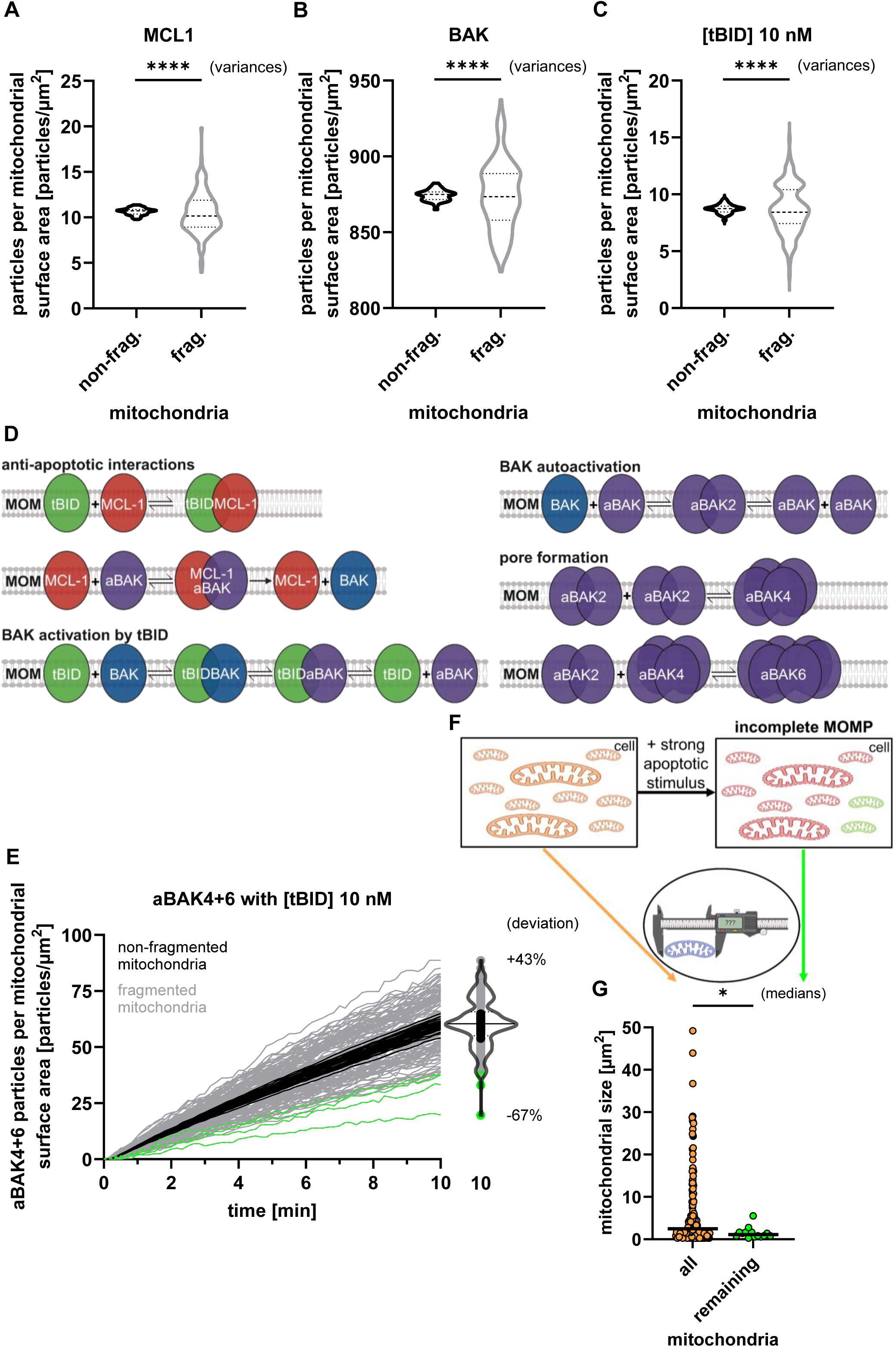
Mitochondrial fragmentation increases variability in MOMP susceptibility due to stochasticity in protein distributions. (A-C) Quantification of protein densities for MCL-1 (A), BAK (B), and 10 nM tBID (C), at the steady-state (0 min) prior to simulating protein interactions. Medians comparison by Man-Whitney U test (p > 0.05). Variances comparison by Levene’s test (p < 0.0001). (D) Schematic of the implemented mitochondrial interaction network showing anti-apoptotic MCL-1 (red), pro-apoptotic tBID (green), and BAK in inactive (blue) and active (purple) states. (E) Formation of BAK pore complexes (aBAK4+6) was simulated over time with 10 nM tBID. Each line represents a single mitochondrion. Results from fragmented (grey) and non-fragmented (black) mitochondria are shown. End-point distributions are visualized as violin and scatter plots. Fragmented mitochondria, most likely to remain intact, are highlighted in green. (F) Schematic hypothesis: Only a subset of fragmented mitochondria remains intact after strong apoptotic stimulation (majority MOMP). (G) Quantification of mitochondrial surface areas from movie S4 of (17). “All” refers to the full population at timepoint 00:00 h, “remaining” to mitochondria not undergoing MOMP. Medians comparison by Mann-Whitney U test (p = 0.0343); n = 176 (all), n = 14 (remaining).

When triggering our model with the strong tBID stimulus of 10 nM, we could observe that non-fragmented mitochondria behaved highly similarly in accumulating substantial amounts of BAK pore oligomers (aBAK4+6) (**Fig. 4E**, black). In contrast, fragmented mitochondria displayed significantly higher variance in their amounts of BAK pore complexes, with a number of fragmented mitochondria presenting only low amounts of BAK pore oligomers (**Fig. 4E**, green). In living cells, a sufficiently high apoptotic stimulus leads to highly synchronized and typically complete MOMP across all mitochondria (13,15). However, if post-mitochondrial apoptosis execution is blocked and cells are provided with sufficient energy, e.g. through glycolytic ATP production, individual mitochondria that evade MOMP might be observed (17,43). Our results on the stochastic heterogeneity in mitochondrial MOMP sensitivity, as most pronounced in fragmented mitochondria, would suggest that especially small mitochondria might be more prone to escape MOMP induction. To test this, we quantified the sizes of mitochondria that either underwent or evaded MOMP in MCF-7 cells expressing Smac-GFP (Movie S1 from (17)) (**Fig. 4F**). We found that mitochondria resisting MOMP were among the smallest in the population (**Fig. 4G**). This observation is consistent with our model predictions and supports a role for stochasticity in shaping MOMP outcomes.

### Stochasticity contributes to fractions of fragmented mitochondria exhibiting high MOMP sensitivity under low apoptotic stress

We next also simulated a condition of low apoptotic stress by reducing the tBID concentration to 2 nM. Fragmented mitochondria exhibited a high variance in mitochondrial-bound tBID per surface area when compared to non-fragmented mitochondria (**Fig. 5A**). It is noteworthy that this distribution is skewed by the distribution hitting the lower limit of zero particles, whereas the high particles densities on fragmented mitochondria were free to rise far higher than those on large, non-fragmented mitochondria (**Fig. 5A**).

**Figure 5:**
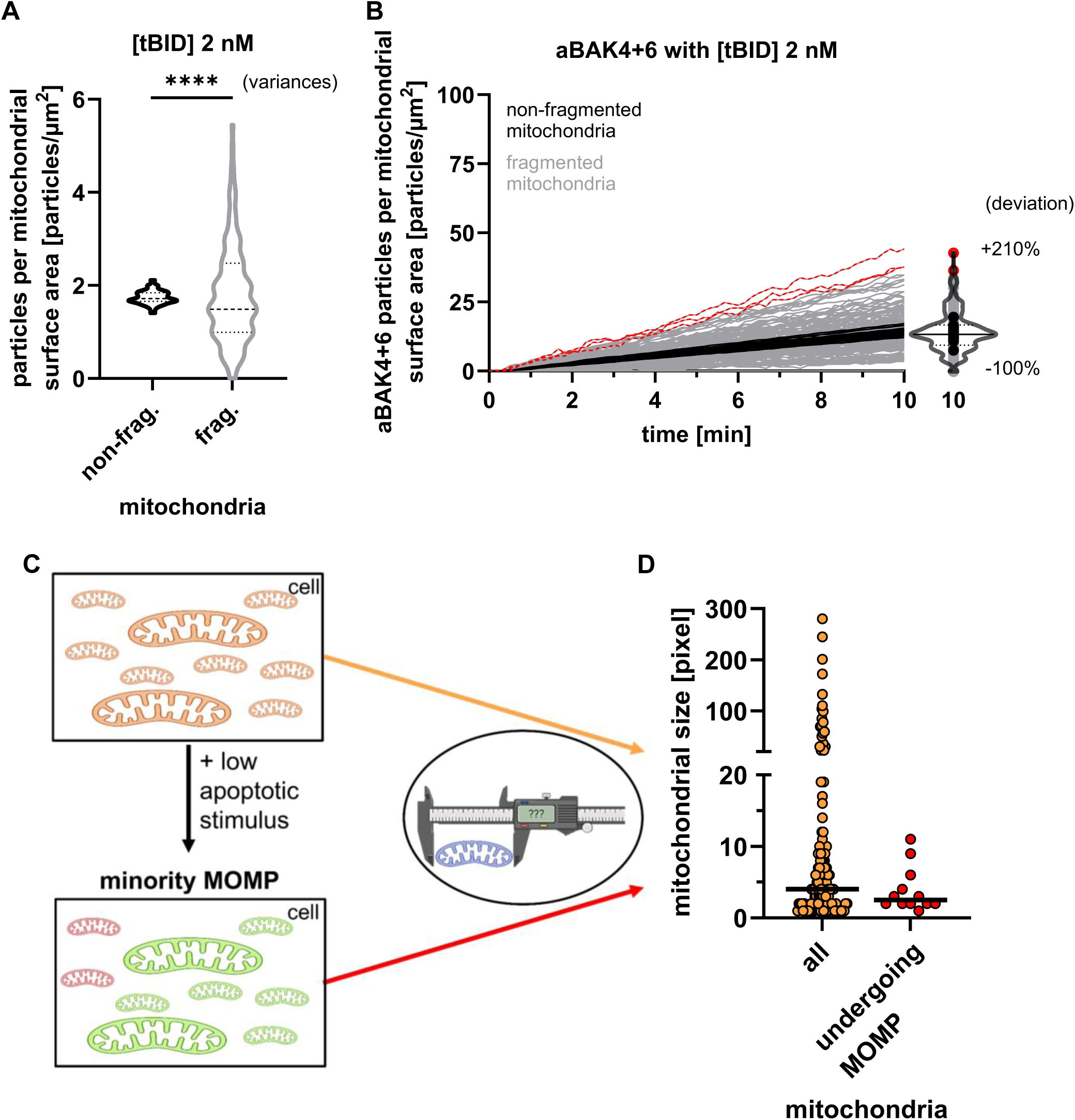
Stochasticity contributes to fractions of fragmented mitochondria exhibiting high MOMP sensitivity under low apoptotic stress. (A) Steady-state quantification of 2 nM tBID associated with mitochondria before simulating protein interactions. Medians comparison by Mann-Whitney U test (p > 0.05). Variances comparison by Levene’s test (p = 6.87e-08). (B) Simulation of aBAK4+6 complex formation over time using 2 nM tBID. Each trace represents a single mitochondrion. Final distributions of activated mitochondria are visualized using violin-scatter plot. Mitochondria most likely to undergo MOMP are highlighted in red. (C) Schematic hypothesis: A subset of fragmented mitochondria undergoes MOMP after a low apoptotic stimulus (minority MOMP). (D) Mitochondrial sizes (in pixels) were measured from Figure S1F/G of (16), comparing the full population with mitochondria undergoing MOMP. Medians are displayed. Medians comparison by Mann-Whitney U test (p = 0.1435); n = 125 (all), n = 12 (undergoing MOMP).

Simulations of the full interactome revealed that the variance in activated BAK oligomers (aBAK4+6) per mitochondrial surface area remained fairly stable between non-fragmented mitochondria (**Fig. 5B**, black). A subset of fragmented mitochondria, however, accumulated substantially more BAK pore complexes than the mean (**Fig. 5B**, red), and these mitochondria would therefore be expected to preferentially undergo MOMP.

While MOMP is traditionally understood as a binary (yes/no) and global mitochondrial response (13), more recent work has demonstrated that sublethal apoptotic stress can induce minority MOMP, a phenotype where MOMP occurs in only a fraction of the mitochondrial population (16,19). Since our simulation results suggest that preferentially small or fragmented mitochondria should undergo MOMP upon sublethal apoptotic stress, we measured mitochondrial sizes in U2OS cells in which only fractions of mitochondria underwent MOMP (Fig. S1F,G of (16)). Quantitative analysis revealed that mitochondria undergoing MOMP in this setting were smaller than the average of the overall mitochondria pool (**Fig. 5D**). This finding is therefore in line with our predictions that stochasticity promotes sub-pools of small mitochondria to preferentially undergo MOMP under conditions of low apoptotic stress.

## Discussion

Our study reveals that stochastic variability plays a causal role in shaping heterogeneity in MOMP susceptibility among mitochondria within individual cells, particularly affecting smaller mitochondria. Stochastic effects in our particle-based model explain how some small mitochondria can resist strong apoptotic signals, while others undergo MOMP even under mild apoptotic stress. Experimental validation supported these predictions: (1) smaller mitochondria were more likely to persist after majority MOMP, and (2) mitochondria undergoing minority MOMP tended to be among the smallest in the population. These findings suggest that mitochondrial fragmentation amplifies stochastic effects in the distribution and activation of apoptosis regulators, thereby enhancing heterogeneity in mitochondrial fates.

Deterministic mathematical models of MOMP have provided valuable insight into the regulation of mitochondrial apoptosis signalling and death decision making (14,39,44,45) but intrinsically cannot accommodate the reaction and movement probabilities required to estimate the influence of stochasticity. We therefore approached this question using particle-based modelling, wherein we deliberately followed a step-by-step model building and expansion strategy that allows to assess if and how stochasticity most likely emanates at subcellular and inter-mitochondrial scale. A key finding from the fundamental model, which includes MCL-1 diffusion within the cytosol and mitochondrial membranes, as well as its exchange between these compartments, was that inter-mitochondrial heterogeneity in MCL-1 amounts increases when mitochondria are distributed irregularly throughout the modeled cell, rather than being evenly spaced. This heterogeneity, observed for a single reactant in the absence of binding partners, likely arises from variations in the available space between mitochondria, being more restricted in densely packed regions and more expansive where mitochondria are sparser. Such spatial organization is consistent with living cells, where mitochondria are typically more concentrated around the nucleus (35,46). In the simplified cell geometry used in our model, this effect is likely underestimated, as we omit several factors that could locally modulate free diffusion. These include diffusion barriers such as the nucleus and other organelles, variations in organelle packing density, as well as constraints on lateral diffusion within mitochondrial membranes.

In addition to mitochondrial positioning and density, our results also demonstrate that mitochondrial fission and fragmentation are strong contributors to inter-mitochondrial heterogeneity in MOMP susceptibility. This finding highlights that, at the scale of small mitochondria and under physiologically relevant protein concentrations, such sources of heterogeneity are significant and should not be overlooked. Mitochondrial fusion and fission are dynamic, ongoing processes in cells, with their balance modulated by environmental conditions and regulated by intracellular signaling pathways (47). For example, mitochondrial remodelling acts as a cellular quality control mechanism to maintain homeostasis by selectively eliminating damaged mitochondria (37,38,48,49). Fusion and fission dynamics are tightly regulated and respond to various cellular stress conditions. Under scenarios such as starvation (50) or infections (51,52) mitochondria undergo morphological adaptations to buffer and resolve stress signals.

The increase in mitochondrial heterogeneity following fission may not be limited to MOMP sensitivity. For instance, studies of mitochondrial fusion and fission dynamics have shown that fission events are often asymmetric in terms of mitochondrial bioenergetics, as evident by mitochondrial membrane potentials of daughter mitochondria. The more polarized daughter mitochondrion has a higher likelihood for subsequent fusion, while the less polarized one is more preferentially cleared by mitophagy (53).

Within the MCL-1/tBID/BAK interactome, our simulations reveal that mitochondrial fission and fragmentation give rise to heterogeneous subpopulations, some more susceptible to apoptotic permeabilization, others exhibiting increased resistance to MOMP. This dual effect helps reconcile the seemingly contradictory roles of fragmented mitochondria: contributing both to recovery after majority MOMP and to sublethal damage during minority MOMP (16,17,19,43). In the context of majority MOMP, the distribution of BCL-2 family proteins across mitochondria has not been thoroughly analyzed. However, in minority MOMP, evidence suggests that pore-forming and anti-apoptotic proteins are unevenly distributed among mitochondria that underwent or resisted MOMP (19). Our particle-based modeling approach suggests that stochastic effects substantially contribute to this and might differentially prime, especially small mitochondria to MOMP.

While stochasticity is not the only factor determining mitochondrial MOMP susceptibility, it is arguably the least accessible through experimental approaches. Multiple factors have been experimentally shown to modulate MOMP susceptibility. For instance, the lipid composition of the mitochondrial outer membrane affects pore formation, with certain lipids such as sphingolipids influencing the activity and oligomerization of pore-forming proteins (20–22). Additionally, membrane curvature defined by the presence and distribution of cardiolipin affects the ease of pore formation (23,24). These biochemical and biophysical features thus can contribute to heterogeneous MOMP susceptibilities across mitochondria. Mathematical modeling, when combined with realistic, experimentally derived parameters as in this study, can therefore complement experimental approaches by revealing how stochastic molecular dynamics influence mitochondrial fate decisions under apoptotic stress.

Stochastic modeling enables the explicit tracking of individual molecular interactions in space and time, a level of detail not attainable with traditional compartment-based, deterministic approaches. The simulation of molecular diffusion and interactions at molecular resolution, incorporating stochastic behavior, has a well-established history (54–56). However, the computational burden of mapping and simulating each individual reactant over time is substantial and difficult to scale. With our implementation of a three-reactant cell-size model, we reached the memory limits of graphic processing unit (GPU)-based high-performance computing. Although the BCL-2 family interactome is more complex than what we could represent in our model, we were still able to qualitatively demonstrate that stochasticity is a key driver of MOMP susceptibility in both majority and minority MOMP scenarios that so far has been underappreciated.

## Materials and methods

### Requirements of the particle-based model

All the codes used to build the particle-based model were created and developed using the FLAME-Accelerated Signalling Tool (FaST version 1.3, downloadable from: https://doi.org/10.5281/zenodo.3757764) (57). FaST-generated codes for graphic processing unit (GPU)-accelerated particle-based models were produced using MATLAB (version 2021a), using the previously described workflow (58). The particle-based models were produced using the FLAME GPU software development kit (SDK) version 1.5 (https://flamegpu.com/download/) and compiled using CUDA 10.2. All simulations were performed on NVIDIA A100 GPUs provided through the BWUniCluster 2.0 (https://wiki.bwhpc.de/e/BwUniCluster2.0), a linux-based cluster running Red Hat Enterprise Linux (RHEL) 8.4.

### Modifications to FaST-generated simulation codes

FaST was used to generate a simplified model of the possible protein interactions using the workflow described previously (58). Input files required for FaST were generated based on the data found in supplemental figures 1A, 1E and 1F. The FaST-generated GPU-based particle-based model simulation codes were implemented and modified as follows. The initial simulation states, defined by FaST in the *OCreatar.c* file, was adapted to include mitochondria as agents. The mitochondria were implemented as rectangular shaped and they did not change their localization over time within a simulation. The model cell’s size was implemented using periodic boundary conditions, defined in the *glabals.h* file, along with other global parameters. The parameters of agents and reactions were specified in the *AgentVariables.h* and *ReactianVariables.h* files. Functions were implemented in the *functians.c* file, where interactions of cytoplasmic or mitochondrial-bound proteins and complexes (e.g, MCL-1_cyto_, MCL-1_mito_ or tBID-MCL-1 complex) with each other was defined by FaST and interactions with mitochondrial agents manually added. Specifically, modifications were made to the “Ligand Output” functions, allowing cytoplasmic proteins (in FaST named as ‘ligands’) to either reflect off or bind to mitochondria after a collision, based on a defined translocation probability. Once bound to the mitochondrial membrane, the protein was reclassified as a mitochondrial membrane-bound protein (in FaST named as ‘receptor’). Conversely, in the “Receptor Output” functions, mitochondrial membrane-bound proteins were allowed to unbind with a defined probability, reverting to cytoplasmic proteins (ligands). Additionally, mitochondrial membrane-bound proteins were enabled to diffuse along the mitochondrial surface. The *XMLMadelFile.xml* file that incorporates all framework parameters, including agents, state transitions, messaging, and simulation layers, was modified to include mitochondria as a separate agent.

### Experimentally-driven parameterization of the particle-based model

A detailed description of the parameter definition, estimation strategies, and validation procedures underlying the particle-based simulation of MCL-1 translocation and mitochondrial protein–protein interactions is provided in the Supplementary Information. These include the stepwise development of simulation parameters, comparative analyses with experimental data, and the implementation of a global parameter estimation approach to constrain the model behavior within physiologically relevant bounds.

### Development of a particle-based model for simulating protein reactions at single-mitochondrion resolution

Simulating MOMP susceptibility on all mitochondria of a representative cell was computationally too expensive due to the number of reactants and species to be simultaneously modelled. We therefore conducted separate simulations of all individual mitochondria per cell to then pool the results into a depiction of the entire pool of mitochondria. The particle distributions and amounts were taken from the final time point (30 minutes) of the whole-cell simulation. Locations on the corresponding mitochondria were then randomly seeded based on the original amounts.

Thereby, a previously established ODE model (39) was modified and parameterized using parameter estimation as described in detail within the Supplementary Methods. This model, incorporating the reaction network and kinetic parameters (**Supplemental Fig. 1E**, **F**), served as the foundation for defining the particle-based mitochondrial model using FaST. The estimated kinetic parameters were based on whole-cell data. To adapt the forward reaction rates k_est_ to single-mitochondrion resolution, these rates were scaled according to the mitochondrial volume (V_m_) to cell volume (V_c_) ratio:

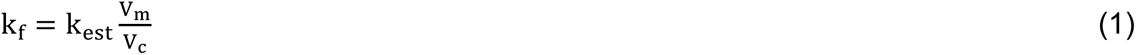

The adjusted second order and first order reaction rates derived from Equation 1 and parameter estimation were incorporated into FaST to compute binding radii, unbinding radii, and unbinding probabilities for the particle-based model (**Supplemental Fig. 1E**, **F**). The *functians.c* and *XMLMadelFile.xml* files from the previously developed particle-based model were modified by removing ligand-related functions and deactivating retro-translocation of particles. Retro-translocation was disabled because the number of mitochondria was sufficient to capture a representative snapshot of mitochondrial heterogeneity, including potential protein retro-translocations. The concentration of BAK was estimated by comparative densitometry analysis of Western blots of NCI-H460 cells.

### Visualization of particle distributions on mitochondrial surfaces

To assess the spatial distribution of particles on mitochondrial membranes and ensure that the implementation of mitochondrial fragmentation did not introduce unintended artifacts, heatmaps of particle intensities were generated. These visualizations were created in Python 3.11 using Matplotlib and NumPy, with additional support from PyVista for 3D rendering. XML files containing particle data were parsed using the integrated XML parser. Heatmaps displayed particle concentrations as intensity values projected onto the surface plane of individual mitochondria. The heatmaps confirmed that no abnormal clustering or edge effects occurred in either fragmented or non-fragmented mitochondrial populations (**Supplemental Fig. 4**). Notably, intensity differences were observed between the two mitochondrial classes, likely reflecting the more uniform spatial distribution of particles across the larger surface areas of non-fragmented mitochondria.

### Measurement of mitochondria sizes from fluorescence imaging data

Images for validation in Figure 4G were obtained from movie S4 of (17) and images for validation in Figure 5D were obtained from Figure S1F/G of (16). The fluorescence microscopy images were analysed in ImageJ (version 1.54g) by converting them to 8-bit format (“Image > Type > 8-bit”) and applying a threshold (“Image > Adjust > Threshold”). If a scale bar was present in the images, the scale was set accordingly. Mitochondrial sizes were quantified by measuring particle sizes using the “Analyze > Analyze Particles…” function.

### Simulation replicates

Simulation replicates of the particle-based model were run where indicated. The particle and/or mitochondrial spatial distributions of the initial files were different. This means that the 0Creator.c file was always used to create a new unique initial state file with a different distribution of particles or mitochondria in the model cell by the seeding of the random number generator with the time. Importantly, the random number generator in FLAME GPU is hard coded with a constant seed, so that all simulations with the same initial states will reproduce the same simulation results.

### Statistics

Tests for normality were performed using GraphPad Prism 10.4.1. All other statistical tests were done in Python 3.11 using the following packages: pandas, SciPy, scikit-posthocs, NumPy. D’Agostino-Pearson normality tests indicated a mixture of normal and non-normal distributions for our data. To ensure consistency in statistical analysis, non-parametric tests were used throughout the paper for comparison of groups (Mann-Whitney U test [single comparison] or Kruskal-Wallis with Dunn’s correction [multiple comparisons]). For comparison of variances, Levene’s test was used for single comparisons and Brown-Forsythe test was used for multiple comparisons.

### Visualisation of reaction networks

Schematics of reaction networks were created with BioRender.com (Fig. 1A, 4D, 4F and 5C).

### Code availability

All codes developed for the particle-based models and visualizations are available at: https://doi.org/10.5281/zenodo.15288516.

## Acknowledgements

The authors acknowledge support by the state of Baden-W0rttemberg through bwHPC.

## Author contributions

JG: data acquisition, analysis and interpretation, draft writing; FK: data acquisition, analysis, draft revision; NP: data acquisition, draft revision, supervision; GF: data interpretation, draft writing, supervision; MR: data interpretation, draft writing, supervision, funding acquisition.

## Ethics

Ethics approval was not required for this work.

## Data availability

Data are available from the authors; all codes are deposited publicly as described above.

