## Supplemental information for "Stochasticity contributes to explaining minority and majority MOMP during apoptosis"

### Supplemental materials and methods

#### Defining simulation parameters for particle-based modeling of MCL-1 translocation

Various parameters, including protein concentration, cellular, and mitochondrial characteristics, needed to be quantified and incorporated into the particle-based model. These parameters are described in the following sections.

The implementation of protein concentration as particle numbers was achieved by deriving median MCL-1 concentrations from gated flow cytometry data using a modified version of MAPIT (1). The median MCL-1 concentration was determined to be 101 nM. The BAK concentration in NCI-H460 cells was measured at 1002 nM via comparative densitometry analysis of Western blots. These concentrations were converted into absolute molecule counts using the following formula:

$$\mathrm{N} = \mathrm{c} * \mathrm{V_c} * \mathrm{N_A} \qquad (1)$$

where $\mathrm{N}$ is particle number, $\mathrm{c}$ is the concentration $\left[\frac{\mathrm{mol}}{\mathrm{L}}\right]$, $\mathrm{V}_c$ is the cell volume $[\mathrm{L}]$ and $\mathrm{N_A}$ is Avogadro constant. For this conversion, an in-house measured cell volume of $3.917*10^{-12}$ L ($G_1$ phase) of NCI-H460 cells was used. The calculated particle numbers for the proteins incorporated into the model are provided in **Supp. Fig. 1A**.

The diffusion coefficient for soluble proteins is required by FaST to simulate the random walk (2). Size dependent cellular diffusion coefficients, $\mathrm{D}$, were estimated using the Stokes-Einstein formula and the measured cellular diffusion rate of GFP, as described previously (3):

$$\mathrm{D} = \mathrm{D_{GFP}} * \left(\frac{\mathrm{m_{GFP}}}{\mathrm{m_{MCL-1}}}\right)^{\frac{1}{3}} = 24\ \frac{\mathrm{\mu m^2}}{\mathrm{s}} * \left(\frac{27\ \mathrm{kDa}}{37\ \mathrm{kDa}}\right)^{\frac{1}{3}} = 21.607\ \frac{\mathrm{\mu m^2}}{\mathrm{s}} \qquad (2)$$

where $\mathrm{m_{MCL-1}}$ represents the molecular weight of MCL-1, $\mathrm{m_{GFP}}$ is the molecular weight of GFP and $\mathrm{D_{GFP}}$ is the measured diffusion rate of GFP in cells (4).

A diffusion coefficient of 21.607 $\mu m^2 s^{-1}$ was assigned for MCL-1 and remained unchanged across different model conditions. The diffusion coefficient for a BCL-2-family protein, BCL-XL, bound to lipid bilayers in giant unilamellar vesicles (GUVs) was previously experimentally determined as 4.8 $\mu m^2 s^{-1}$ (5), this value was used as the diffusion coefficient for all BCL-2-family proteins on membranes based on the assumption that molecular weight will have a lower impact on diffusion within a membrane according to the Saffman-Delbrück theory (6).

**Specification of the *in-silico* cell geometry based on high-resolution 3D imaging**

In-house measurements of NCI-H460 geminin cells size from 3D-imaging data were used to construct a representative three-dimensional simplified model cell containing mitochondria. In brief, the acquisition protocol was as follows: untreated NCI-H460 geminin cells were stained with Mitotracker and for MCL-1 and microscopy images were acquired using Airyscan Joint Deconvolution in Superresolution mode. The whole cell volume was measured using the MCL-1 signal across the whole cell. Mitochondrial network parameters (mitochondrial number, mitochondrion surface area and volume) were extracted using the Mitochondria Analyzer pipeline in ImageJ and are shown in **Supp. Fig. 1B**.

This data was used to determine the dimensions of the model cell to ensure an equivalent volume. The mitochondria within the model were assigned to a rectangular shape with the same number, surface area and volume as the analysed mitochondria. The final model parameters for the mitochondria and the cell for the model cell environment are summarized in **Supp. Fig. 1C**.

**Determination of kinetic parameters for MCL-1 translocation**

To implement translocation of MCL-1, fluorescence recovery after photobleaching (FRAP) was combined with steady state measurements of MCL-1 localisation to capture both dynamics (from FRAP) and distributions. FRAP was performed by bleaching MCL-1 on stage, which was endogenously fluorescence tagged via mScarlet knock-in. After the bleaching, the fluorescence recovery within the bleached area was measured, to estimate the translocation kinetics of MCL-1 (**Supp. Fig. 2D**). The FRAP profile was used to obtain the fluorescence recovery rate of 0.00452 $s^{-1}$ using non-linear one-phase association regression analysis (**Supp. Fig. 2E**). In-house imaging data of immunofluorescence-stained MCL-1 combined with markers for mitochondria and nuclei was used to quantify steady state MCL-1 concentrations in the cytoplasm ($MCL\text{-}1_{Cyto}$) and mitochondria ($MCL\text{-}1_{Mito}$) revealing that the $[MCL\text{-}1_{Cyto}]$:$[MCL\text{-}1_{Mito}]$ ratio is 0.7923 (**Fig. 1C**). From these values, a fitting function was derived by integrating the fluorescence recovery rate with the $[MCL\text{-}1_{Cyto}]$:$[MCL\text{-}1_{Mito}]$ ratio, resulting in the function:

$$\mathrm{MCL}-1_{\mathrm{Mito}}[\mathrm{particles}] * \left(1-\mathrm{e}^{\mathrm{k}*\mathrm{t}}\right) = 28671 * \left(1-\mathrm{e}^{-0.00452 s^{-1} * t}\right) \quad (3)$$

Where 28671 represents the absolute amount of MCL-1 particles on the mitochondria to give the expected $[MCL\text{-}1_{Cyto}]$:$[MCL\text{-}1_{Mito}]$ ratio of 0.7923. This data was used to estimate $k_{on}$ and $k_{off}$ values of MCL-1 translocation dynamics. A least-squares global parameter estimation was conducted using a basic ODE model:

$$\frac{\mathrm{d}[\mathrm{MCL}-1_{\mathrm{Cyto}}]}{\mathrm{dt}} = \mathrm{k}_{\mathrm{off}}[\mathrm{MCL}-1_{\mathrm{Mito}}] - \mathrm{k}_{\mathrm{on}}[\mathrm{MCL}-1_{\mathrm{Cyto}}] \quad (4)$$

$$\frac{\mathrm{d}[\mathrm{MCL}-1_{\mathrm{Mito}}]}{\mathrm{dt}} = \mathrm{k}_{\mathrm{on}}[\mathrm{MCL}-1_{\mathrm{Cyto}}] - \mathrm{k}_{\mathrm{off}}[\mathrm{MCL}-1_{\mathrm{Mito}}] \quad (5)$$

Estimated upper and lower bounds for $k_{on}$ and $k_{off}$ were set between 0.01 $s^{-1}$ and 0.00001 $s^{-1}$. The values with the lowest estimation cost were selected for further analysis, resulting in $k_{on}$ 0.00054367 $s^{-1}$ and $k_{off}$ 0.0039562 $s^{-1}$. These values were validated by reintroducing them into the basic ODE model, which successfully reproduced the experimentally obtained [MCL-$1_{Cyto}$]:[MCL-$1_{Mito}$] ratio of 0.7923, with the model yielding a ratio of 0.793. These estimated $k_{on}$ and $k_{off}$ values were integrated into the particle-based model by calculating probability values. For translocation of MCL-$1_{Cyto}$ to the mitochondria, first the collision rate of MCL-1 colliding with mitochondria was determined by simulating MCL-1 diffusion in the cytoplasm and calculating collisions with mitochondria. Collisions refers to when a molecule of MCL-$1_{Cyto}$ crosses a reflective boundary of a mitochondrion. The collision rate, of 195.169862 $s^{-1}$ and the $k_{on}$ of 0.00054367 $s^{-1}$ estimated above were used to define the binding probability:

$$\text{probability}_{\text{binding}} = \frac{k_{on}}{\text{collision rate}} = \frac{0.00054367\ s^{-1}}{175.169862\ s^{-1}} = 3.10367 * 10^{-6} \quad (6)$$

Likewise, the estimated $k_{off}$ value was used to calculate the unbinding probability, describing the MCL-$1_{Mito}$ detachment from the mitochondrial membrane:

$$\text{probability}_{\text{unbinding}} = 1 - e^{-k_{off} * \Delta t} = 1 - e^{-0.0039562\ s^{-1} * 0.05\ s} = 1.9779 * 10^{-4} \quad (7)$$

A function was implemented to generate a random number between 0 and 1 to determine if binding upon collision or unbinding occurs after each timestep. In the event that a collision of MCL-$1_{Cyto}$ did not lead to a binding event, the MCL-$1_{Cyto}$ particle was reflected off the mitochondrial surface. In the event of binding, the MCL-1 molecule was assigned as MCL-$1_{Mito}$ and localised at the point of intersection with the mitochondrial surface.

**Parameter estimation for simulating mitochondrial protein–protein interactions**

Previously, we published an ODE-model of BCL-2-family interactions, incorporating a representative of the BH3-only proteins (tBID), an anti-apoptotic protein (BCL-XL) and a pore-forming effector protein (BAX) (7). This model, specifically its reaction network, was used as the basis for the particle-based models of events at individual mitochondria. We, however, made the following changes. Firstly, we used MCL-1 instead of BCL-XL as a representative anti-apoptotic protein on the basis that though functionally similar, we have recently demonstrated experimental capture of MCL-1 localization behavior in cells during the cell cycle (8). This data and the experimental tools generated from this study and ongoing follow up studies allow us to better experimentally define MCL-1 steady state and retrotranslocation dynamics than for BCL-XL. Importantly, key kinetic rates, such as retrotranslocation of BAX and BAK, are consistent between BCL-XL and MCL-1 (9). Secondly, we used BAK instead of BAX as a representative pore-forming effector protein. Whilst functionally similar, BAX is predominantly found in the cytoplasm but migrates to the mitochondria upon apoptotic stress whereas BAK is predominantly located at the mitochondrial membrane. The use of BAK allowed us to neglect the effect of dynamic exchange of BAX between mitochondria and the cytoplasm, which is highly computationally challenging, and focus on capturing snapshots of MOMP-sensitivity. Importantly, as (retro)translocation times are generally slower than diffusion dynamics, the effect of stochasticity on BAX distributions is expected to be generally higher than for BAK. Therefore, despite these changes, our findings are relevant to the extended network of BCL-2-family interactions.

To parameterize the model, it was necessary to undertake a parameter estimation to obtain a single parameter set that represents both the expected dynamic behavior of BCL-2-family interactions and reproduces quantitative experimental reference data. This section describes the estimation of these parameters.

**Mitochondrial Model Parameter Estimation: Definition of Reference Data**

To undertake the parameter estimation, 2 sets of experimental reference data were extracted from the publication of Subburaj and colleagues (Fig. 3G and 4D from (10)) using ImageJ. The first data set describes the formation of BAX oligomers for 1 h in the presence of cBID on supported lipid bilayers (SLBs) mimicking the composition of the mitochondrial membrane (hereafter referred to as Estimation Data 1). The second data set describes the BCL-XL-mediated disruption of pre-formed BAX oligomers after 1 h (hereafter referred to as Estimation Data 2). In both experiments, the readout was percentage occurrence of different BAX oligomers on SLBs (monomers, dimers, trimers, tetramers, pentamers and hexamers). Notably, the experimental setup used for Estimation Data 2 includes a 1 h pre-formation of BAX oligomers by incubation of BAX and cBID with the exact same experimental conditions as used to generate Estimation Data 1. Therefore, a parameter estimation was performed based on 2 sequential simulations representing both Estimation Data 1 and 2. First, a 1 h simulation was performed using the initial conditions shown in **Supp. Fig. 1D**.

At the end of the first simulation (Simulation 1), the cost (Cost 1) was calculated from the end points of the simulation and Estimation Data 1:

$$\mathrm{Cost\ 1} = \sum(\mathrm{model} - \mathrm{experiment})^2 \quad (8)$$

A detailed explanation of how model data and experimental data were prepared and compared is given in the next section. Thereafter, the end of Simulation 1 was used to initialise Simulation 2 with all species and intermediates from Simulation 1 after 1 h carried over with the addition of 2.5 nM of BCL-XL. Simulation 2 was then performed for 1 h and the cost for Simulation 2 calculated (Cost 2) in the same way as Cost 1 (Equation 8). The total cost for a parameter set was given by:

$$\mathrm{Total\ Cost} = \mathrm{Cost\ 1} + \mathrm{Cost\ 2} \quad (9)$$

**Mitochondrial Model Parameter Estimation: Comparing Simulation and Experimental Values**

The experimental data was measured as percentage occurrence of each BAX oligomer on the SLB membrane after 1 h. As no trimers and pentamers were observed experimentally and were explicitly excluded from the model reaction network, the model and experimental levels of monomeric, dimeric, tetrameric and hexameric BAX were used for the cost analysis. For monomers of BAX from simulations, the quantities of aBAX, tBID-bound BAX and BCL-XL-bound BAX were added together to represent all species of BAX expected to be detectable within membranes (Equation 10). For dimers, tetramers and hexamers, the concentration of the equivalent species was taken directly from the model (aBAX2, aBAX4 and aBAX6 respectively). As the experimental data was based on area under the curve measurements from the fitting of six Gaussian curves (representing monomers, dimers, trimers, tetramers, pentamers and hexamers) to a frequency distribution (fluorescence intensity vs number of events) to estimate percentage occurrence of each species, the concentration of BAX multimers from simulations was multiplied by the oligomerisation state (i.e. 2× for dimers, 4× for tetramers). To normalise the simulation data to percentage occurrence, the sum of all BAX species, including oligomers, expected to be detectable within membranes (i.e. all except inactive BAX monomers) multiplied by their respective BAX oligomerisation state was calculated (see Equations 10-13).

$$\text{BAX monomer occurrence }(\%) =$$

$$100\% \cdot \frac{[\text{aBAX}]+[\text{tBIDBAX}]+[\text{tBIDaBAX}]+[\text{BCLXLaBAX}]}{[\text{aBAX}]+2\cdot[\text{aBAX2}]+4\cdot[\text{aBAX4}]+6\cdot[\text{aBAX6}]+[\text{tBIDBAX}]+[\text{tBIDaBAX}]+[\text{BCLXLaBAX}]} \qquad (10)$$

$$\text{BAX dimer occurrence }(\%) =$$

$$100\% \cdot \frac{2\cdot[\text{aBAX2}]}{[\text{aBAX}]+2\cdot[\text{aBAX2}]+4\cdot[\text{aBAX4}]+6\cdot[\text{aBAX6}]+[\text{tBIDBAX}]+[\text{tBIDaBAX}]+[\text{BCLXLaBAX}]} \qquad (11)$$

$$\text{BAX tetramer occurrence (\%)} = 100\% \cdot \frac{4\cdot[\text{aBAX4}]}{[\text{aBAX}]+2\cdot[\text{aBAX2}]+4\cdot[\text{aBAX4}]+6\cdot[\text{aBAX6}]+[\text{tBIDBAX}]+[\text{tBIDaBAX}]+[\text{BCLXLaBAX}]} \quad (12)$$

$$\text{BAX hexamer occurrence (\%)} = 100\% \cdot \frac{6\cdot[\text{aBAX6}]}{[\text{aBAX}]+2\cdot[\text{aBAX2}]+4\cdot[\text{aBAX4}]+6\cdot[\text{aBAX6}]+[\text{tBIDBAX}]+[\text{tBIDaBAX}]+[\text{BCLXLaBAX}]} \quad (13)$$

**Mitochondrial Model Parameter Estimation: Definition of Upper and Lower Bounds**

Model parameters were constrained to biological plausible lower and upper limits similar to our previously developed model (7). For forward reactions, reactions were constrained between 1E3 – 1E6 $M^{-1}\cdot s^{-1}$. $K_D$ ranges of 1E-10 –1E-6 M was previously determined to be reasonable ranges for BCL-2-family interactions (7). Therefore, reverse reaction rates were set by estimating the forward reaction rates and $K_D$s, through the relationship:

$$k_r = k_f \cdot K_D \quad (14)$$

First order enzymatic reactions (**Supp. Fig. 1F**), were constrained between 0.1–1E-7 $s^{-1}$ and retrotranslocation rates (**Supp. Fig. 1E**) were constrained between 0.1–1E-5 $s^{-1}$ as performed previously based on experimentally measured retrotranslocation rates (7).

Initial parameter estimations based on these boundaries yielded an unsatisfactory cost and reproduction of the reference data. Moreover, several parameters were frequently located towards the edge of their constraints. It was therefore reasoned that a neglected aspect of using an ODE paradigm for the parameter estimation was the concentrating effect of several reactions occurring exclusively at mitochondrial membranes (a factor intrinsically addressed in our particle-based simulations). Therefore, for second order reactions that occur exclusively at membranes, the upper bounds for forward rates were increased by 100× (reflecting an ~10-fold concentrating effect on both reactants). Reciprocally, the $K_D$ lower bounds for these reactions were lowered by 100×. The resulting parameter estimation constraints (see **Supp. Fig. 1D**) led to a vastly improved reproduction of the reference data.

**Mitochondrial Model Parameter Estimation: Implementation of a Global Parameter Estimation**

The parameter estimation was performed in MATLAB (R2020b) using previously established scripts for a global parameter estimation (2). In brief, initial parameter sets were selected within pre-defined parameter ranges by Latin hypercube sampling. For each initial parameter set, a local minima was determined using the MATLAB function fmincon (https://de.math-works.com/help/optim/ug/fmincon.html). The parameter estimation was performed in parallel across 12 worker nodes for 72 h, giving a final total of 6153 iterations with local minima. The parameter set with the least global cost were chosen as the best fit parameter set.

**Mitochondrial Model Parameter Estimation: Results**

The global minima from the parameter estimation had a cost of 199.5 and showed reasonable replication of Estimation Data 1 (**Supp. Fig. 2A**) and Estimation Data 2 (**Supp. Fig. 2B**). Full simulation outputs showed a strong effect of the addition of BCL-XL in the rapid de-assembly of BAX oligomers and retrotranslocation of BAX. This strong inhibitory effect of anti-apoptotic BCL-2 family proteins on pore formation and particularly the role played by retrotranslocation was shown by us to be an integral feature of apoptotic signalling to achieve the generally all-or-nothing response of mitochondrial outer membrane permeabilization (7). Based on this and its good reproduction of quantitative reference data, these parameters were then used in the particle-based simulations of mitochondrial pore formation.

### Figure legends

**Supp. Figure 1: Experimental data and parameterization for constructing a particle-based model of a mitochondrial network in a model cell.**

(A) Protein concentrations of selected BCL-2 family members in NCI-H460 cells.

(B) Mean morphological parameters derived from untreated NCI-H460 geminin cells used for constructing a simplified model cell.

(C) Mitochondrial and cellular dimensions in the model cell were defined to match the experimentally derived measurements shown in (B). Mitochondria were represented as rectangular compartments with dimensions approximating the experimentally observed shapes.

(D) Initial concentrations for simulation of BAX-oligomer formation. Initial values were taken from (10).

(E) Protein interactions and corresponding reaction parameters of second order reactions. The full parameterization strategy is described in detail in the Methods section.

(F) Protein interactions and corresponding reaction parameters of forward reactions. The full parameterization strategy is described in detail in the Methods section.

**Supp. Figure 2: (A-C) Replication of experimental kinetics and dynamics of BAX oligomerization.**

(A)-(C) Comparison of model and experimental data for Estimation Data 1 (A) and Estimation Data 2 (B). Experimental datapoints were extracted from Figure 3G (A) and 4D (B) in (Subburaj et al., 2015).

(C) Time-course dynamics of BAX oligomerization representing Experiment 1 (before addition of BCL-XL) and Experiment 2 (after addition of BCL-XL).

**Sup. Figure 2: (D-E) Experimental kinetic data for subcellular localization of MCL-1.**

(D) Normalized mean fluorescence intensities from FRAP experiments with MCL-1.

(E) Fluorescence recovery was fitted using a non-linear one-phase association model, yielding a rate constant of 0.00452 $s^{-1}$. Details of how this data were used to define MCL-1 localization dynamics in the model are provided in the Methods section.

**Supp. Figure 3: Control simulations and model parameters for mitochondrial fragmentation.**

(A) Overview of model cell and mitochondrial parameters used for simulations involving mitochondrial fragmentation (see Fig. 3A). Cell size and total mitochondrial volume were kept constant across all conditions.

(B) Second independent simulation run of Fig. 3D. Median values did not differ significantly (Kruskal-Wallis, $p > 0.05$), but variance was consistently higher in fragmented mitochondria (Brown-Forsythe, $p < 0.0001$ across all concentrations).

(C) Third independent simulation run of Fig. 3D. As in (B), variance was significantly increased in fragmented mitochondria at all concentrations (Brown-Forsythe, $p < 0.0001$), while median values remained unchanged (Kruskal-Wallis, $p > 0.05$).

**Supp. Figure 4: Visualization of homogenous particle distributions across mitochondrial surface planes.**

Particle distributions were visualized using Matplotlib in combination with NumPy, displaying particle concentrations as intensity values across the mitochondrial surface plane. To analyse spatial distributions, each mitochondrial surface was divided into bins of 10 nm in length to capture local concentration differences with high spatial resolution.

**Supplemental Fig. 1**

**A**

| Protein | Concentration (nM) | Particle Number |
|---|---|---|
| MCL-1 | 101 | 237307 |
| BAK | 1002 | 2365498 |
| tBID | 1 | 2359 |
| | 2 | 4718 |
| | 5 | 11795 |
| | 10 | 23591 |
| | 20 | 47182 |
| | 50 | 117956 |

**B**

**Mean values of NCI-H460 cell measurements**

| Parameter | |
|---|---|
| Mitochondria number | 45 |
| Mitochondrion surface area | 59 $\mu m^2$ |
| Mitochondrion volume | 8 $\mu m^3$ |
| Cell volume | 3917 $\mu m^3$ |

**C**

| | Mitochondrion | Cell |
|---|---|---|
| length [nm] | 580.003 | 10000 |
| height [nm] | 580.003 | 4500 |
| width [nm] | 25428.1 | 87053.777 |
| volume [$nm^3$] | 8.554E+09 | 3.91742E+12 |
| surface area [$nm^2$] | 5.966E+07 | - |

**D**

| Protein | Initial Concentration (nM) |
|---|---|
| tBID | 5 |
| BAX | 2.5 |
| BCL-XL | 0 |

**E**

| | Reactions | $k_f$ ($M^{-1}s^{-1}$) | $k_r$ ($s^{-1}$) | PBM Binding Distance (nm) | PBM Unbinding Distance (nm) | PBM Dissociation Probability |
|---|---|---|---|---|---|---|
| 1 | BAK + aBAK ↔ aBAK2 | 372661 | 4.04E-04 | 0.82 | 3.28 | 2.01E-06 |
| 2 | aBAK + aBAK ↔ aBAK2 | 5.51E+07 | 2.12E-03 | 9.99 | 39.96 | 1.05E-05 |
| 3 | aBAK2 + aBAK2 ↔ aBAK4 | 9.99E+07 | 3.54E-02 | 13.45 | 53.80 | 1.77E-04 |
| 4 | aBAK2 + aBAK4 ↔ aBAK6 | 1.4E+07 | 7.2E-03 | 5.04 | 20.17 | 3.60E-05 |
| 5 | tBID + BAK ↔ tBIDBAK | 204027 | 3.73E-02 | 0.60 | 2.43 | 1.86E-04 |
| 8 | tBID + aBAK ↔ tBIDaBAK | 3.7E+07 | 1.79E+01 | 8.18 | 32.73 | 8.54E-02 |
| 9 | tBID + MCL1 ↔ tBIDMCL1 | 4.8E+07 | 4.17 | 9.32 | 37.30 | 2.06E-02 |
| 10 | MCL1 + aBAK ↔ MCL1aBAK | 7.83E+07 | 3.02E-01 | 11.90 | 47.63 | 1.50E-03 |

**F**

| | Forward Reactions | $k_f$ ($s^{-1}$) | PBM Dissociation Probability |
|---|---|---|---|
| 6 | tBIDBAK → tBIDaBAK | 0.099 | 4.98E-04 |
| 7 | tBIDaBAK → tBIDBAK | 1.23E-07 | 6.13E-10 |
| 11 | MCL1aBAK → MCL1 + BAK | 0.1 | 4.99E-04 |

Supplemental
Fig. 2

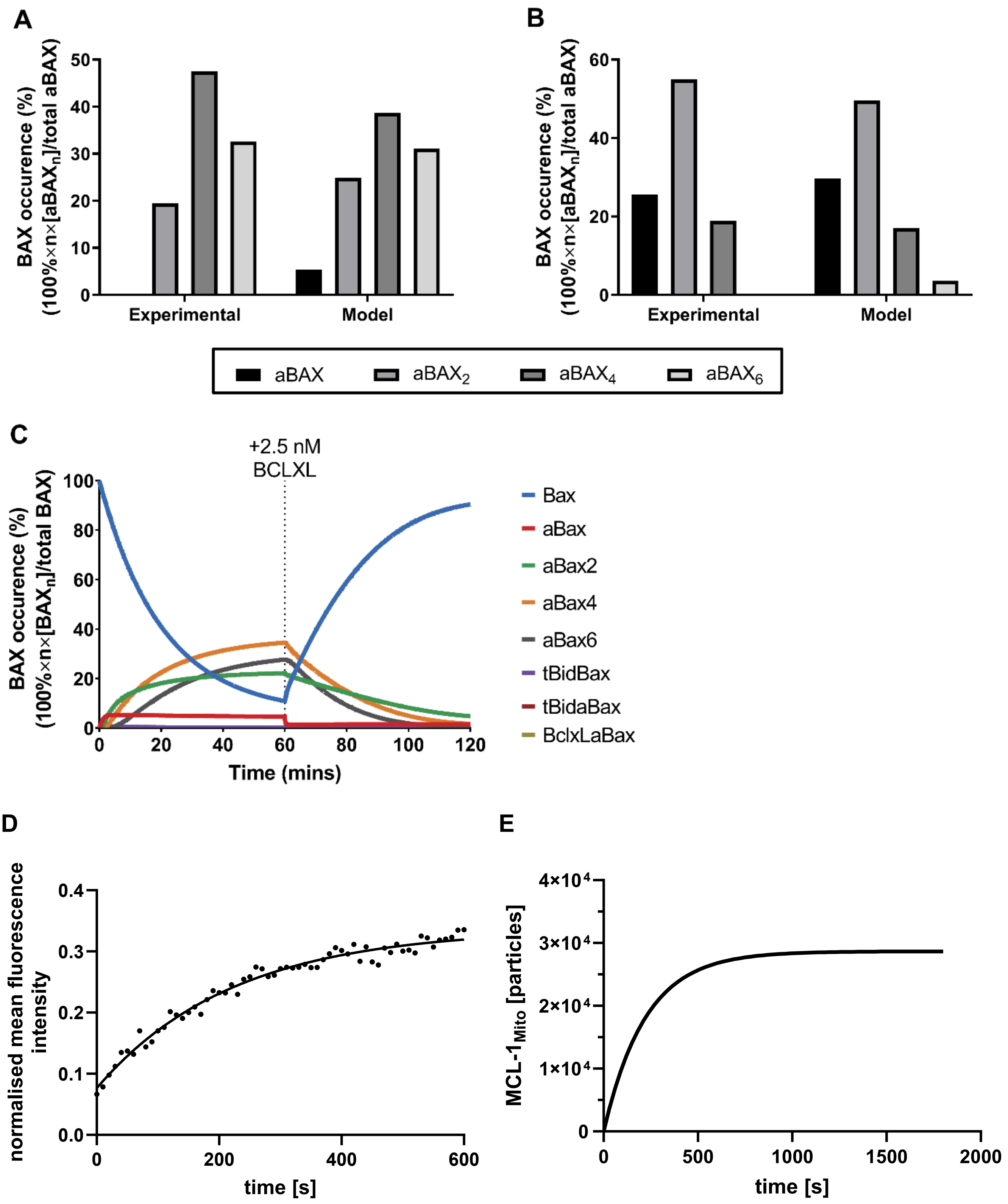

**Supplemental Fig. 3**

**A**

| Parameter | Model variant: only non-fragmented mitochondria | Model variant: non-fragmented and fragmented mitochondria | |
|---|---|---|---|
| Mitochondria number | 45 | 40 | 158 |
| Mitochondrion surface area | 59 μm$^2$ | 59 μm$^2$ | 2 μm$^2$ |
| Mitochondrion volume | 8 μm$^3$ | 8 μm$^3$ | 0.2 μm$^3$ |
| Cell volume | 3917 μm$^3$ | 3917 μm$^3$ | |
| MCL-1 concentration | 101 nM | 101 nM | |
| MCL-1 particle number | 237 307 | 237 307 | |

**B**

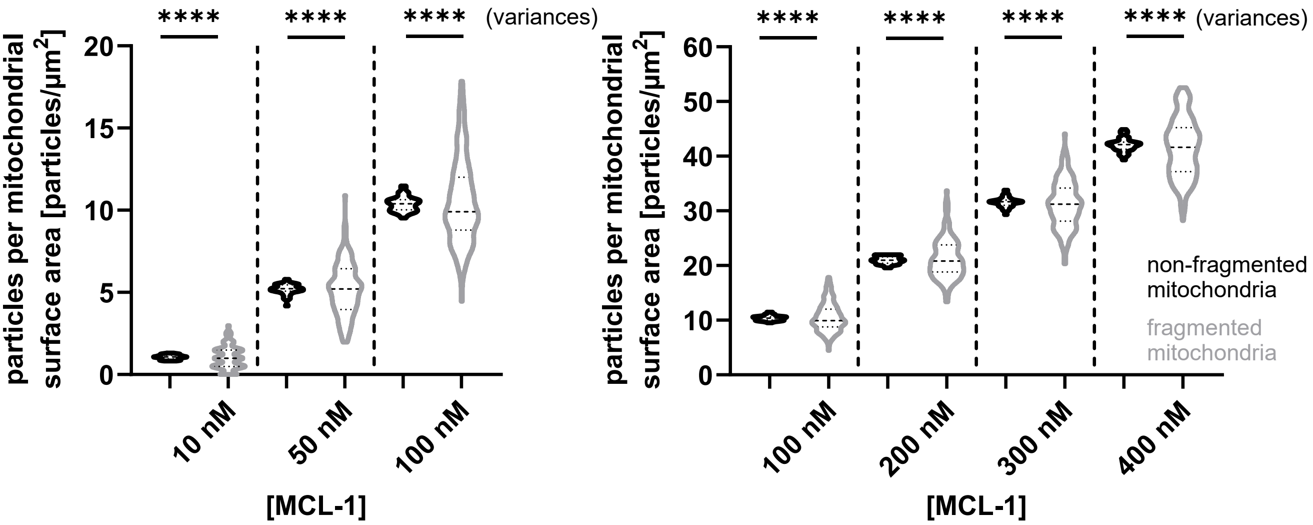


**C**

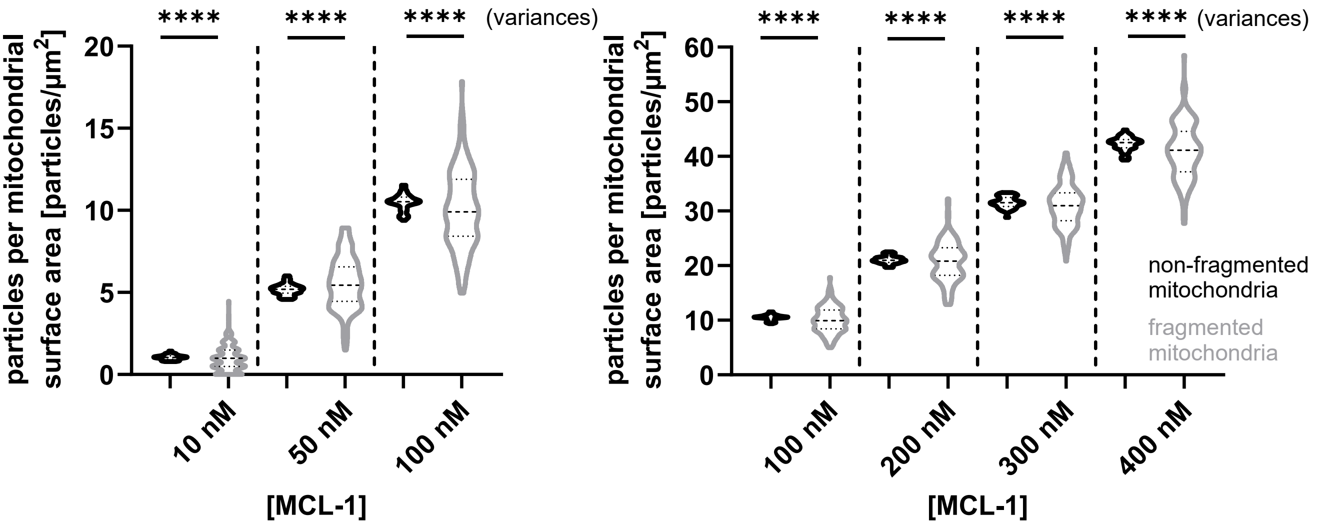

Supplemental Fig. 4

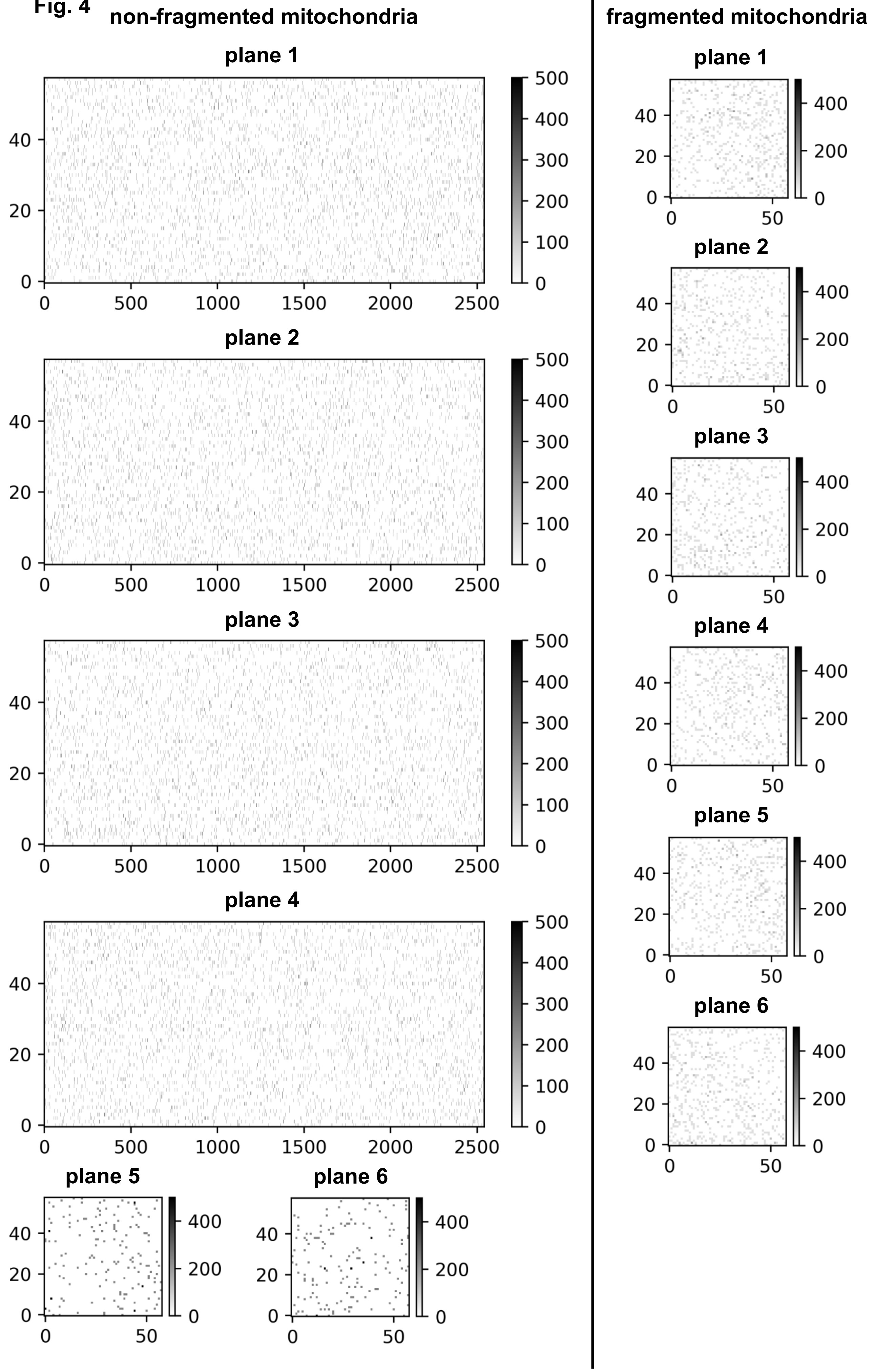
